# Machine-learning guided Venom Induced Dermonecrosis Analysis tooL: VIDAL

**DOI:** 10.1101/2023.05.21.541619

**Authors:** William Laprade, Keirah E. Bartlett, Charlotte R. Christensen, Taline D. Kazandjian, Rohit N. Patel, Edouard Crittenden, Charlotte A. Dawson, Marjan Mansourvar, Darian S. Wolff, Thomas J. Fryer, Andreas H. Laustsen, Nicholas R. Casewell, José María Gutiérrez, Steven R. Hall, Timothy P. Jenkins

**Affiliations:** Department of Applied Mathematics and Computer Science, Technical University of Denmark, Kongens Lyngby, Denmark; Centre for Snakebite Research and Interventions, Liverpool School of Tropical Medicine, Liverpool, UK; Department of Biotechnology and Biomedicine, Technical University of Denmark, Kongens Lyngby, Denmark; Instituto Clodomiro Picado, Facultad de Microbiología, Universidad de Costa Rica, San José, Costa Rica

**Keywords:** Dermonecrosis, Snakebite envenoming, Machine learning, VIDAL, Venom, Necrosis, Mouse models, Toxinology, Antivenom, Neglected tropical disease

## Abstract

Snakebite envenoming is a global public health issue that causes significant morbidity and mortality, particularly in low-income regions of the world. The clinical manifestations of envenomings vary depending on the snake’s venom, with paralysis, haemorrhage, and necrosis being the most common and medically relevant effects. To assess the efficacy of antivenoms against dermonecrosis, a preclinical testing approach involves *in vivo* mouse models that mimic local tissue effects of cytotoxic snakebites in humans. However, current methods for assessing necrosis severity are time-consuming and susceptible to human error. To address this, we present the Venom Induced Dermonecrosis Analysis tool (VIDAL), a machine-learning-guided image-based solution that can automatically identify dermonecrotic lesions in mice, adjust for lighting biases, scale the image, extract lesion area and discolouration, and calculate the severity of dermonecrosis. We also introduce a new unit, the dermonecrotic unit (DnU), to better capture the complexity of dermonecrosis severity. Our tool is comparable to the performance of state-of-the-art histopathological analysis, making it an accessible, accurate, and reproducible method for assessing dermonecrosis. Given the urgent need to address the neglected tropical disease that is snakebite, high-throughput technologies such as VIDAL are crucial in developing and validating new and existing therapeutics for this debilitating disease.

## Introduction

Snakebite envenoming is a major public health problem, especially in low-income regions of the world^1^. Indeed, it is responsible for substantial morbidity and mortality, particularly in the impoverished areas of the tropics and subtropics, such as sub-Saharan Africa, South and Southeast Asia, Papua New Guinea, and Latin America^2–4^. Whilst accurate estimates are difficult to obtain, it is believed that between 1.8–2.7 million people worldwide are envenomed each year, resulting in 80,000-140,000 deaths and around 400,000 victims left with permanent sequelae^5–7^. Notably, snake venoms are highly complex and comprise a wide range of toxins that differ across families, genera, and species. Consequently, the clinical manifestations and pathophysiological effects of envenomings can vary greatly depending on which snake species was responsible for the snakebite, with paralysis, haemorrhage, and necrosis some of the most common and/or medically relevant effects^8^. This also results in the need to test different venoms for the preclinical validation of antivenom efficacy. To complicate matters further, the severity of a given envenoming is significantly affected by the amount of venom injected and the anatomical location of the bite^9^. When a new antivenom is developed, or an existing one is introduced to a new geographical setting, it needs to undergo preclinical efficacy testing; this involves the assessment of its neutralising capacity against the lethal effects of venom(s) in mice^10,11^, but it may also involve a diverse set of tests to assess neutralisation of other relevant toxic effects, such as necrosis within the skin (i.e., dermonecrosis). This pathology is predominately caused by cytotoxic 3FTxs (three-finger toxins), SVMPs (snake venom metalloproteinases), and PLA2s (phospholipase A2s), which induce significant tissue damage by disrupting cell membranes, breaking down extracellular matrix, and inducing inflammation. Venom induced dermonecrosis often results in surgical intervention, e.g. debridement or amputation of the affected limb^12–14^, and frequently results in loss of limb function or permanent disability.

The underlying mechanism by which snake venoms induce dermonecrosis, the characterisation of necrotising toxins, and their varying severities have, for a long time, presented a key area of fundamental and translational research within the field of Toxinology. As necrosis typically affects cutaneous and muscle tissues in snakebite victims, these are the two types of tissue most studied using *in vivo* mouse models to mimic the local tissue effects of cytotoxic snakebites in human victims^15^. Necrosis within the muscle tissue (myonecrosis) is primarily assessed by injecting venom or toxins into the gastrocnemius muscle of mice and assessing the necrosis-inducing potential via the quantification of the extent of muscle damage by histological analysis (i.e. haemotoxylin & eosin (H&E)-staining) or by quantifying the plasma activity of creatine kinase (CK)^16^. Alternatively, dermonecrosis caused by snake venoms is tested using methods initially described by Theakston, *et al*., in which mice are injected intradermally with sub-lethal doses of venom (with or without venom-inhibiting treatments) to induce tissue damage within the skin’s layers, and after 72 hours, the mice are euthanised and the width and height of the venom-induced lesions are measured using callipers ^17^. While able to determine if a treatment drastically inhibits the cytotoxic effects of venoms, this method is susceptible to human error, can be inaccurate, and does not take lesion severity into account (*i*.*e*., a light lesion and dark lesion of the same size but with different intensities would be regarded as equally severe using this method). In an attempt to better differentiate lesion severity, one option is to analyse and quantify dermonecrosis severity within each skin layer using histopathological analysis of H&E-stained lesion cross-sections ^18^. However, this method is time consuming, requires analysis and quantification by trained pathologists, and is also susceptible to human error.

Given the current drive towards addressing the neglected tropical disease that is snakebite, major efforts are being undertaken into defining pathologies, understanding which toxins are responsible, as well as testing and developing new and existing therapeutics. Thus, high throughput, accurate, and reproducible technologies are key in ensuring that these efforts are as time- and cost-efficient as possible. Therefore, in this article, we present a new and accessible machine-learning guided solution, i.e the Venom Induced Dermonecrosis Analysis tooL, VIDAL (https://github.com/laprade117/VIDAL). Here, we trained a machine learning algorithm to automatically identify dermonecrotic lesions in mice using photography images, adjust for lighting biases, scale the image, extract lesion area and discolouration, and calculate the severity of dermonecrosis. We also propose a new unit to better capture the complexity of dermonecrosis severity, *i*.*e*., the dermonecrotic unit (DnU). We validate the utility of this tool for quantitatively defining dermonecrosis using samples derived from animal models of envenoming and demonstrate our tool is comparable to the performance of the current state of the art histopathological analysis.

## Results

In this study, we present our VIDAL image analysis tool trained on 193 sample images sourced from diverse experiments (Fig.1). All venoms used induced dermonecrosis in mice, as observed macroscopically, and the intensity of dermonecrosis was dependent upon the snake species from which the venom was collected and the venom dose injected. To illustrate the appropriateness of our methods, we present histopathological analyses for nine example lesions that are subsequently also assessed with VIDAL, *i*.*e*., three healthy tissue controls only injected with PBS (C1_A, C1_B, C2_A); three light lesions caused by West African *N. nigricollis* (57 μg; L1_A, L1_B,) and *N. pallida* (25 μg; L2_A) venom; and three dark lesions caused by East African *N. nigricollis* (63 μg; D1_A, D1_B) and West African *N. nigricollis* (57 μg; D2_A) venom.

**Fig. 1.**
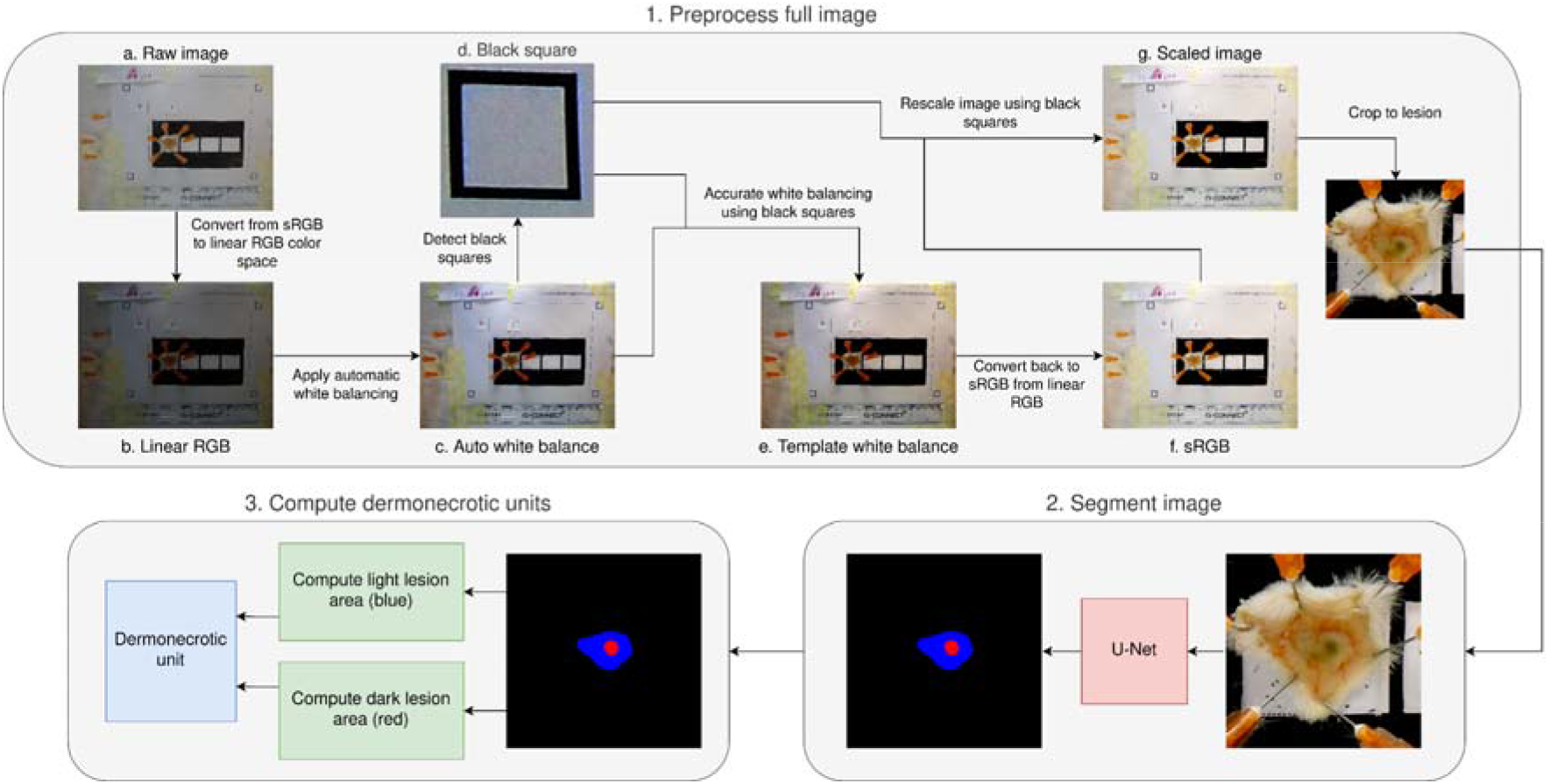
Overview of the workflow for VIDAL. First, the raw image is imported and converted from sRGB to linear RGB. Thereafter, the image is white balanced and subsequently further white balanced using the colour of the paper detected via the scaling squares. In parallel, the image is rescaled using the same squares. This processed image is then used for segmentation and automatic identification of the necrotic lesions. Together, this information is used to compute the dermonecrotic lesion area and differentiate between 1. a severe dark dermonecrotic lesion (red) and 2. a less severe light region of local tissue damage (blue). The areas of each dermonecrotic and light region of local tissue damage are then combined based on our histopathologically determined (c.f. 2.4) weighting of 2.019 to 1 into dermonecrotic units (DnU).

### Histopathological analysis of H&E-stained sections of venom-induced lesions demonstrate clear differences in light and dark lesions

Skin tissue samples from mice injected with PBS showed the characteristic histological features of normal skin, including epidermis, dermis, hypodermis, panniculus carnosus, and adventitia layers. In contrast, the skin layers of mice injected with snake venoms showed various degrees of damage, depending on the type and dose of venom. In the case of tissue sections collected from macroscopically dark lesions, the extent of damage was more pronounced than in macroscopically light lesions. Using the dermonecrosis scoring system developed previously for quantifying lesion severity in H&E-stained lesion cross-sections ^19^, the healthy tissue, light lesions, and dark lesions each received mean overall dermonecrosis severity scores of 0.00, 1.73, and 3.50, respectively (Fig. 2, Fig. S1). The dark versus light lesion severity weighting of 2.019 was then calculated by dividing the dark by the light lesion mean overall dermonecrosis severity score.

**Fig. 2.**
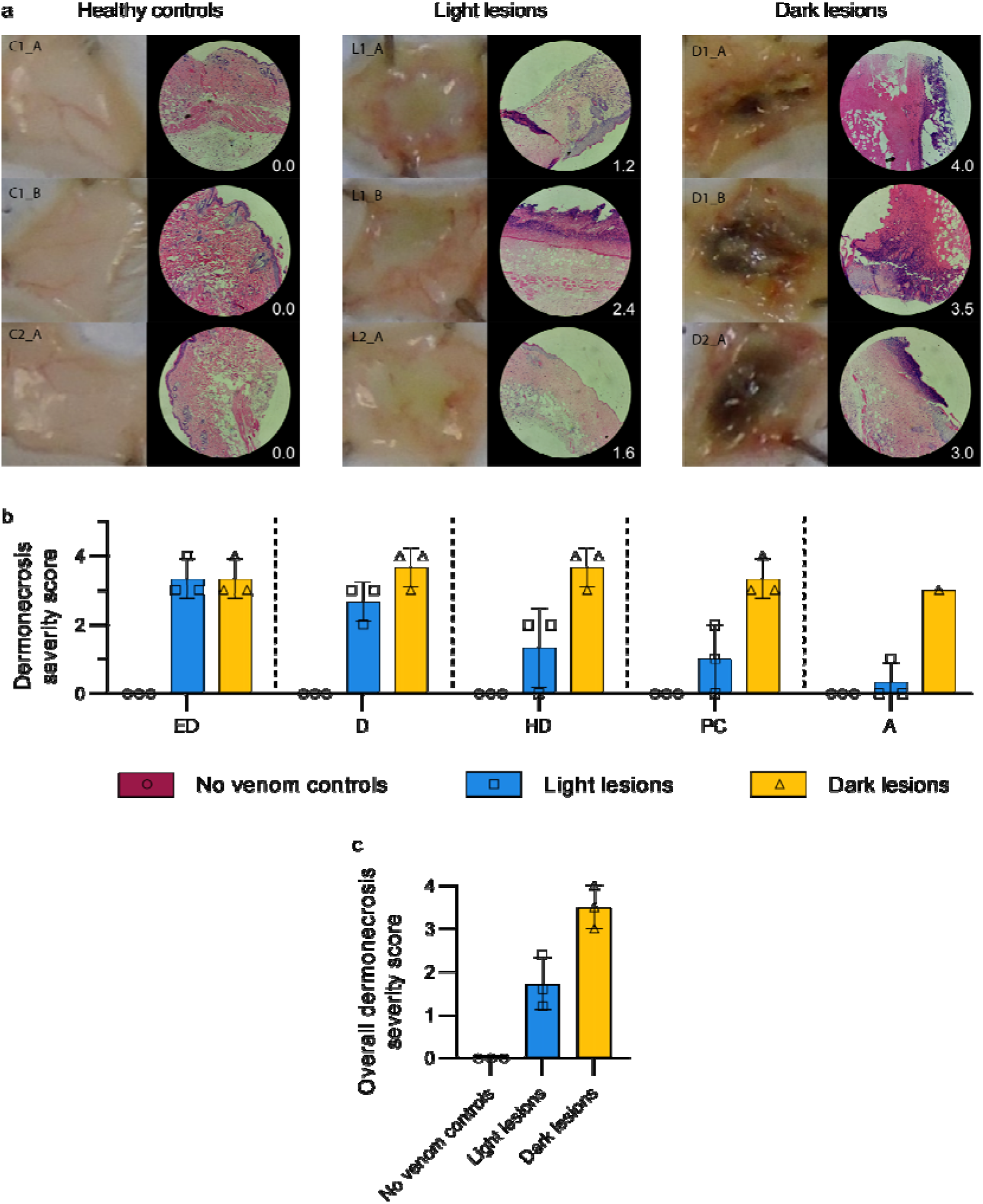
Histopathological analysis of mouse skin cross-sections of healthy tissue, light lesions, and dark lesions confirms dark lesions are more severely dermonecrotic than light lesions. Four μm thick H&E sections from formalin-fixed, paraffin-embedded tissue were previously prepared from the dermal injection sites of mice 72 hours after being ID-injected with PBS (C1_A, C1_B, C2_A) or dermonecrotic snake venoms, *i*.*e*., three light lesions caused by West African *N. nigricollis* (57 μg; L1_A, L1_B,) and *N. pallida* (25 μg; L2_A); three dark lesions caused by East African *N. nigricollis* (63 μg; D1_A, D1_B) and West African *N. nigricollis* (57 μg; D2_A). A pathologist (JMG) scored, between 0-4, the percentage of necrosis that was visible within each skin layer, where 0 = 0%, 2 = 25-50%, 3 = 50-75%, and 4 = 75-100%, based on previously developed methodology ^19^. The highest recorded score per skin layer was taken as the measure of maximum severity it reached, and the mean score of all layers per sample taken as its overall dermonecrosis severity score. (**a**) Macroscopic and representative 100x-magnified H&E-stained images with their associated overall dermonecrosis severity scores in their bottom right-hand corners can be seen for (left) healthy tissue controls (C1_A, C1_B, C2_A), (middle) light lesions (L1_A, L1_B, L2_A), and (right) dark lesions (D1_A, D1_B, D2_A). Bar graphs summarising the (**b**) measured level of dermonecrosis in each skin layer [epidermis (ED), dermis (D), hypodermis (HD), panniculus carnosus (PC), and adventitia (A)], and (**c**) mean overall dermonecrosis severity score for each lesion or control tissue sample. All H&E-stained tissue images used in the measurements can be seen in Fig. S4.

### White balancing is able to normalise for different lighting conditions

The tool automatically applies white balancing to account for potential lighting variations, ensuring consistent results across images. The white balancing function operates effectively, yielding comparable outcomes (Fig. 3).

**Figure 3.**
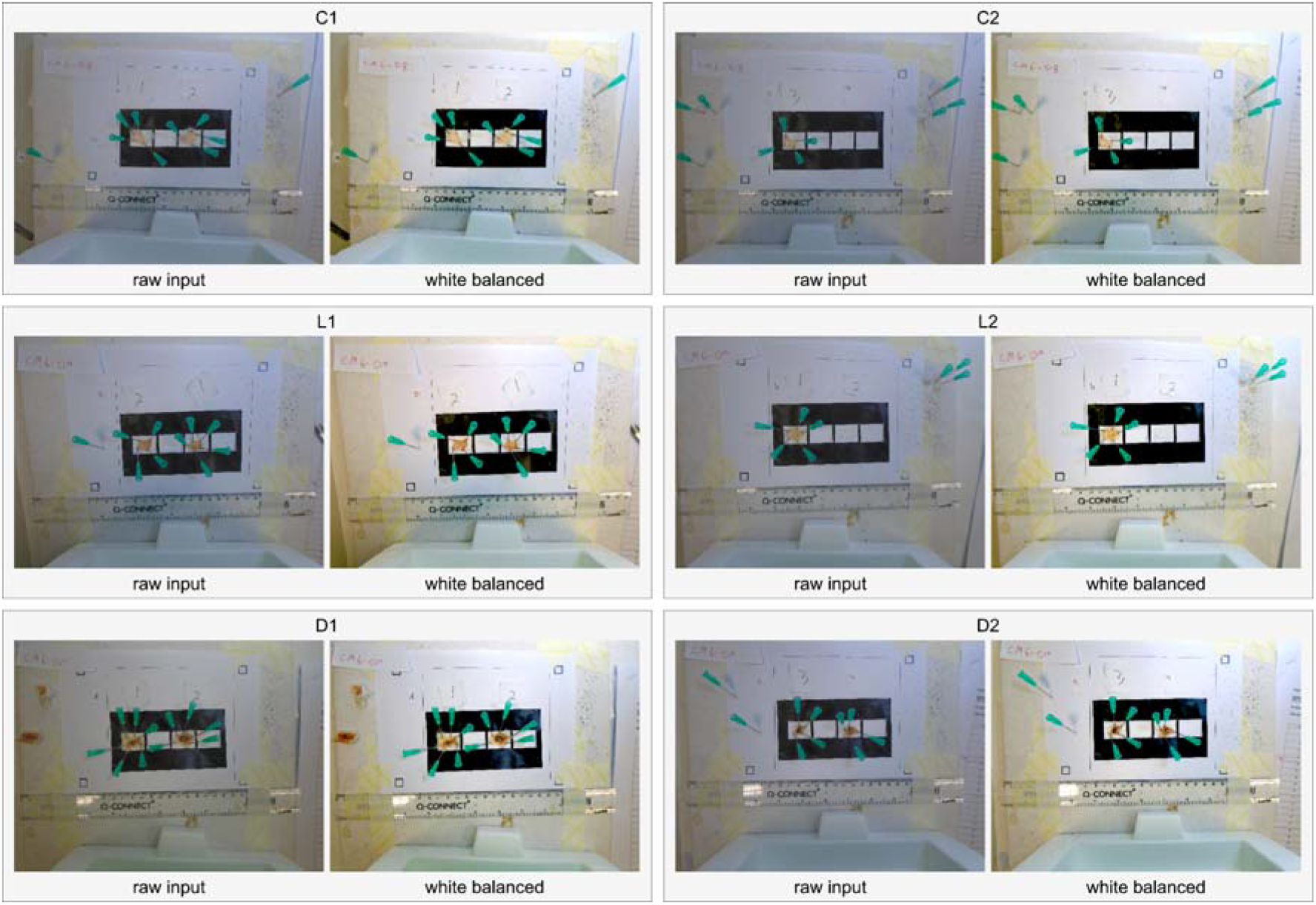
An overview of VIDAL’s automatic white balancing output across the 9 example lesions. Included were three healthy tissue controls only injected with PBS (C1_A, C1_B, C2_A); three light lesions caused by West African *N. nigricollis* (57 μg; L1_A, L1_B,) and *N. pallida* (25 μg; L2_A); three dark lesions caused by East African *N. nigricollis* (63 μg; D1_A, D1_B) and West African *N. nigricollis* (57 μg; D2_A).

In order to conduct a more rigorous evaluation of the white balancing feature’s ability to adapt to challenging lighting conditions, we introduced image manipulations by randomly simulating various colours. In each simulation, the tool successfully restored an image that was similar in quality (Fig. 4).

**Figure 4.**
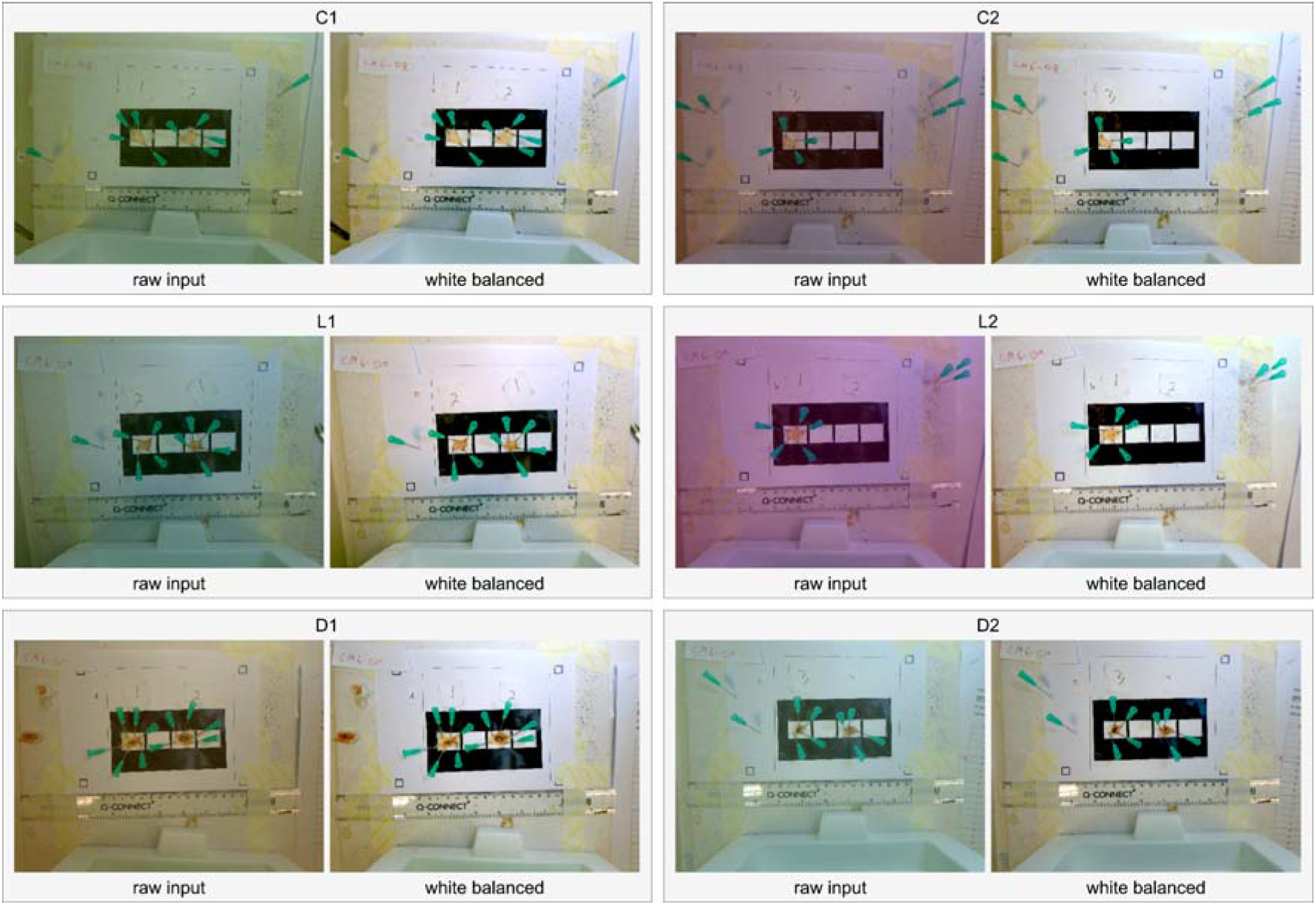
Stress testing VIDAL’s automatic white balancing capacity. The colours of the six images were randomly manipulated to assess white balancing performance. Included were three healthy tissue controls only injected with PBS (C1_A, C1_B, C2_A); three light lesions caused by West African *N. nigricollis* (57 μg; L1_A, L1_B,) and *N. pallida* (25 μg; L2_A); three dark lesions caused by East African *N. nigricollis* (63 μg; D1_A, D1_B) and West African *N. nigricollis* (57 μg; D2_A).

### Scaling is able to ensure scale normalisation of images

To determine scale the tool first uses a standard template matching algorithm to locate the black squares in the corners of the paper template in each image. Using the known scale of these black squares we can then compute the scale of the images (in pixels per mm) and resize the images to a target scale. We use a target of 5 pixels per mm as this allows us to fit the entire lesion nicely into a 256 × 256 patch that we can feed into the U-Net segmentation model directly (Table 1).

**Table 1.** Table outlining the input dimensions, detected scale, target scale, and output dimensions across the 9 example images. Included were three healthy tissue controls only injected with PBS (C1_A, C1_B, C2_A); three light lesions caused by West African *N. nigricollis* (57 μg; L1_A, L1_B,) and *N. pallida* (25 μg; L2_A); three dark lesions caused by East African *N. nigricollis* (63 μg; D1_A, D1_B) and West African *N. nigricollis* (57 μg; D2_A).

|  | C1 | C2 | L1 | L2 | D1 | D2 |
| --- | --- | --- | --- | --- | --- | --- |
| Input dimensions (pixel x pixel) | 3864 x 5152 | 3864 x 5152 | 3864 x 5152 | 3864 x 5152 | 3864 x 5152 | 3864 x 5152 |
| Detected scale (pixels per mm) | 9.9247 | 10.0936 | 12.8996 | 11.0422 | 13.8578 | 10.5688 |
| Target scale (pixels per mm) | 5 | 5 | 5 | 5 | 5 | 5 |
| Output dimensions (pixel x pixel) | 1947 x 2596 | 1914 x 2552 | 1498 x 1997 | 1750 x 2333 | 1394 x 1859 | 1828 x 2437 |

### Segmentation is able to identify and distinguish between light and dark lesions

To automatically identify lesion areas, the tool uses a U-Net segmentation model to segment the lesions located at user-defined positions in the image. Overall, an average MCC score of 0.7644 and an average F1 (Dice) score of 0.8738 were achieved, and we were able to predict 99.98% of the pixels correctly across 25 runs (Fig. 5).

**Figure 5.**
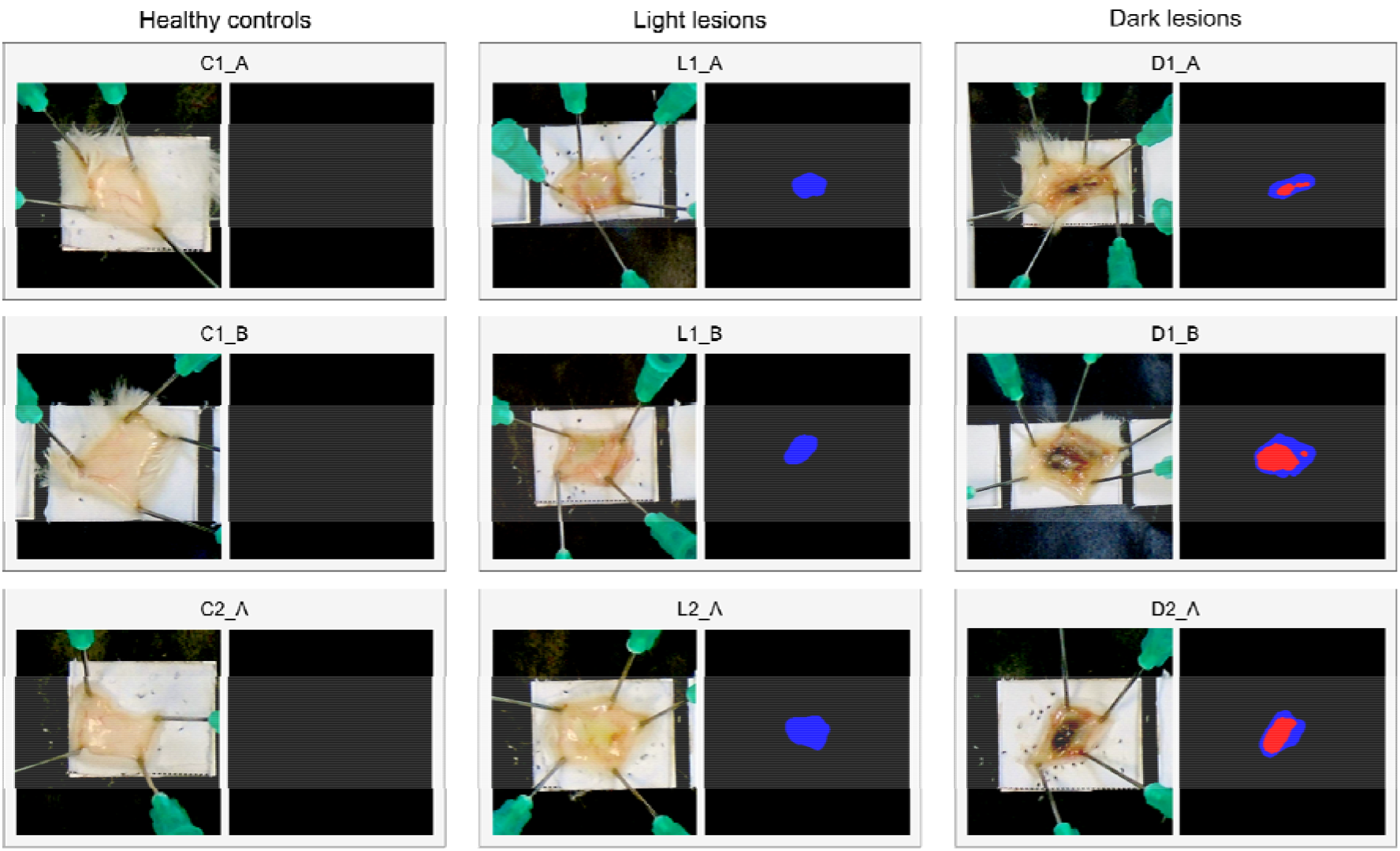
Segmentation of all necrotic lesions across the 9 lesions assessed in the histopathological analysis. The blue regions denote light necrosis and the red regions signify dark necrosis. Included were three healthy tissue controls only injected with PBS (C1_A, C1_B, C2_A); three light lesions caused by West African *N. nigricollis* (57 μg; L1_A, L1_B,) and *N. pallida* (25 μg; L2_A); three dark lesions caused by East African *N. nigricollis* (63 μg; D1_A, D1_B) and West African *N. nigricollis* (57 μg; D2_A).

### Dermonecrotic units present a easy and representative readout for lesion severity

To assess the severity of each lesion, the tool automatically computes the real area and DnU for each mouse skin excision in all of the images (Table 2).

**Table 2.** Dermonecrotic units and lesion sizes for the six example test images. Included were three healthy tissue controls only injected with PBS (C1_A, C1_B, C2_A); three light lesions caused by West African *N. nigricollis* (57 μg; L1_A, L1_B,) and *N. pallida* (25 μg; L2_A); three dark lesions caused by East African *N. nigricollis* (63 μg; D1_A, D1_B) and West African *N. nigricollis* (57 μg; D2_A).

| Results |  |  |  |  |  |  |  |  |  |
| --- | --- | --- | --- | --- | --- | --- | --- | --- | --- |
| Lesion | C1_A | C1_B | C2_A | L1_A | L1_B | L2_A | D1_A | D1_B | D2_A |
| Light area (mm<sup>2</sup>) | 0.00 | 0.00 | 0.00 | 41.96 | 43.28 | 71.00 | 33.08 | 63.04 | 41.24 |
| Dark area (mm<sup>2</sup>) | 0.00 | 0.00 | 0.00 | 0.00 | 0.00 | 0.00 | 12.84 | 55.80 | 44.60 |
| DnU | 0.00 | 0.00 | 0.00 | 41.96 | 43.28 | 71.00 | 58.98 | 175.59 | 131.20 |

### The tool’s graphical user interface is simple and accessible

To ensure accessibility and easy implementation of VIDAL across research, production, and quality control laboratories, a graphical user interface was developed (https://github.com/laprade117/VIDAL). Our tool can be used to quickly upload an image and receive statistics on the lesion area, luminance, and DnU for each mouse in the image (Fig. 6).

**Figure 6.**
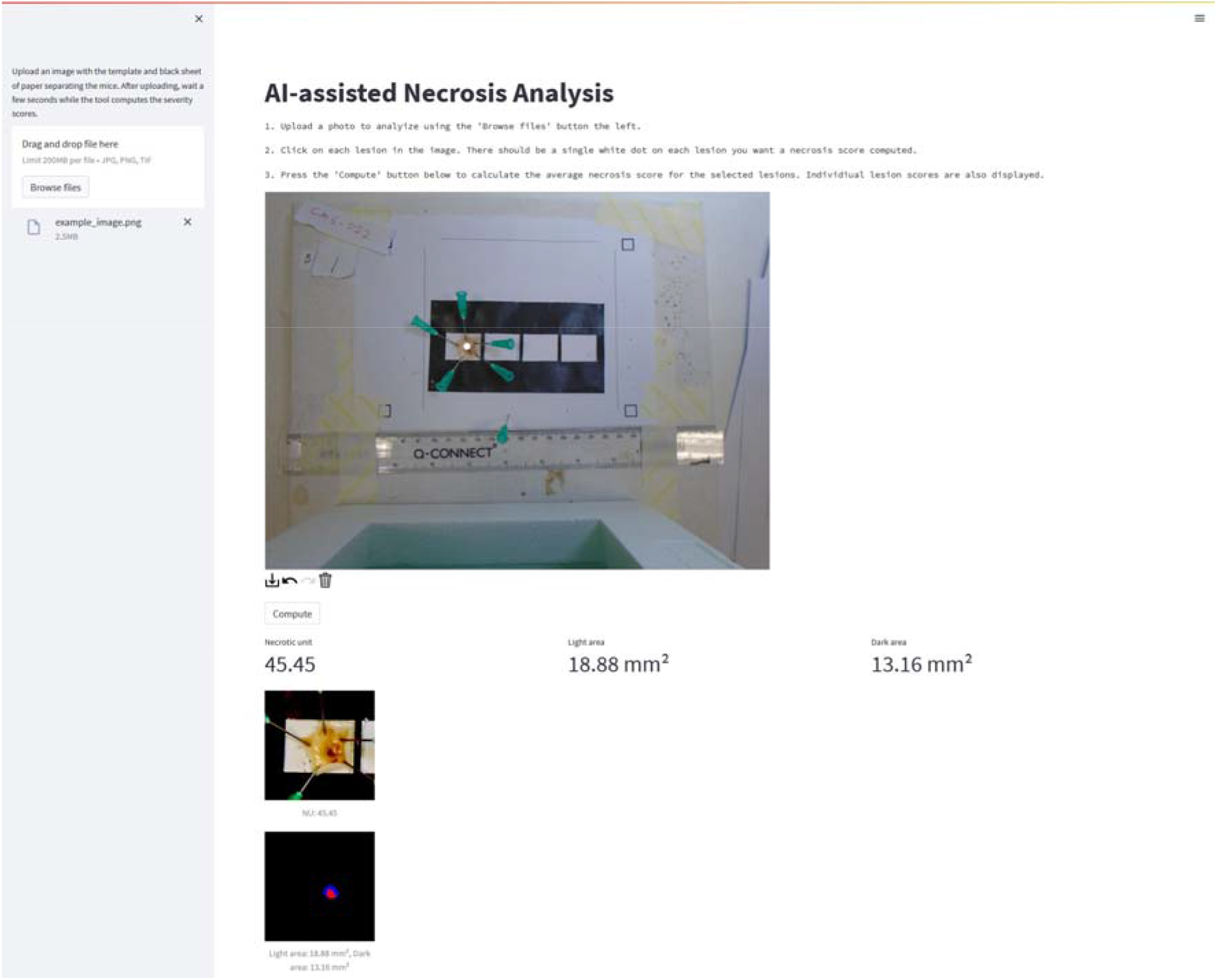
Image of VIDAL’s web interface. By uploading an image file of the experiment, the tool will, within seconds, provide the white balanced reference image, the individual lesions it detected, and how it decided to segment them, as well as all relevant data on lesion areas and DnUs.

## Discussion

Snake venom-induced dermonecrosis presents one of the most severe clinical manifestations of envenoming by viperid and some elapid species and is responsible for a substantial proportion of the estimated 400,000 permanent sequelae induced by snakebite envenomings globally every year^1^. Furthermore, current antivenoms are largely ineffective in preventing the rapid destruction of tissues that snake venoms may cause, unless administered very soon after the bite^1^. Therefore, a thorough understanding of snake venom dermonecrosis and the neutralising potential of current, as well as next generation, therapies for this pathophysiological manifestation is crucial ^12–14,20^. However, to date, no rapid and robust dermonecrosis assays exist that can be used to help create such a thorough understanding. Indeed, current assessments involve the manual measurements of dermonecrotic lesions with callipers ^19–21^ or the use of manual lesion outlining on transparent plastic sheets and calculation of the encompassed area with millimetric paper ^22^. Both of these methods are inherently inaccurate, since they can only assess entirely spherical areas precisely (which dermonecrotic lesions seldomly are), and are susceptible to potential unconscious bias and human error. Additionally, both existing methods entirely fail to evaluate the severity of the dermonecrotic damage, *i*.*e*., a light lesion and dark lesion of the same size would be considered equally severe. This shortcoming can be overcome via the use of histopathological analysis, which is accurate, but low-throughput and requires training as well as specialised equipment ^18,19^.

Until recently, the assessment of snake venom haemorrhage was faced with the same issues. Therefore, we first developed a simple computational tool to improve accuracy of quantifying haemorrhage severity, which has since been implemented by different groups within the field ^23,24,24–31^. This encouraged us to build a more sophisticated machine learning guided tool that further increased the accuracy, accessibility, and reproducibility of snake venom haemorrhage assessment^32^.

In this study, we applied a similar approach to the area of dermonecrosis and developed and evaluated an easy-to-use, rapid, and accurate method for the evaluation of snake venom-induced dermonecrosis in a rodent skin model, which can both measure the exact size of the lesions and provide a severity score based on the presence of light (less severe) versus dark (more severe) lesions. Thereby, a more accurate and rapid measure of dermonecrosis compared to current techniques is achieved. Images generated from envenoming by a variety of viper and elapid venoms that were known to induce dermonecrosis were used to ensure that the results were representative. Histopathological analyses confirmed the different extents of overall skin damage in light versus dark macroscopic lesions, as judged by the more severe pathological alterations observed in various layers of the skin in the latter. We further built a fully automated analysis pipeline, supported by vision AI (U-Net). Our tool, VIDAL, efficiently and accurately evaluated a diverse set of training images that encompassed varying levels of dermonecrotic lesion severity. Throughout our experimentation, we consistently observed reliable white balancing, precise scaling, and accurate segmentation. To further examine the white balancing feature, we artificially manipulated the images to simulate different lighting conditions and colour settings, mimicking scenarios where different cameras were used by different users. Our tool effectively adjusted the white balance of these manipulated images, demonstrating its robustness in experimental settings. To minimize animal usage during the initial validation phase, we performed segmentation on a limited dataset. Nevertheless, U-Net has consistently displayed its robustness in previous studies^33–36^, even when trained on small datasets, which was the case in our study. The segmentation results consistently aligned with expert opinions, even when the images were artificially manipulated to simulate various laboratory and lighting conditions, thus creating a more challenging environment for testing image quality.

In order to ensure maximum accessibility and seamless integration into existing workflows, we have developed a user-friendly web tool with a graphical user interface (GUI). This tool enables users to perform quick and precise analyses of dermonecrotic lesions. By utilizing a smartphone, capturing a photo, uploading the image, and obtaining accurate information regarding the severity of venom-induced dermonecrotic lesions in mice can now be accomplished in a matter of minutes. This remarkable reduction in analysis time from hours to just a few minutes has significantly enhanced efficiency. Additionally, the user-friendly nature of the tool greatly increases accessibility, enabling us to provide a standardized solution that can be utilised in laboratories without extensive training or prior knowledge of lesion assessment requirements. Thereby, this tool may help strengthen the area of toxicovenomics, *i*.*e*., the study of venom proteomes in relation to their functional toxicity^37^, by improving standardisation and harmonisation of toxicity data across venoms, models, and labs. In turn, this may help provide a clearer overview of the key toxin targets that need to be neutralised by antivenoms, and guide the development of novel envenoming therapies^38–40^.

Whilst the tool brings many advantages, it does come with some of the same limitations as our prior tool ^41^. These include potential decreases in white balancing accuracy in very poorly or unevenly lit environments, as well as a heavily damaged template sheet. Additionally, lesions that include significant haemorrhaging are sometimes poorly recognised, potentially complicating the analysis of necrosis induced by some viper venoms; we therefore recommend that users aim to clean such lesions and assess these images on a case-by-case basis to ensure that the program accurately detects their lesions. Finally, though only an issue for certain venoms, the tool could present false positives for some of the very light lesions (*i*.*e*., secondary skin damage) due to similarity to the mouse skin colour; this is primarily an issue in lower resolution images.

## Conclusion

With the Venom Induced Dermonecrosis Analysis tooL VIDAL, we introduce a rapid and robust new method for the automated assessment of venom-induced dermonecrosis in mice by implementing sophisticated machine learning based image analysis approaches. This method eliminates the risk of human biases in assessing lesion areas and increases the speed of analysis substantially. We hope that this will be of utility for the study of dermonecrotic toxins and venoms and provide researchers with an extra tool to be implemented in the assessment of neutralising efficacy of antivenoms and inhibitors.

## Methods

Images of murine dermonecrotic lesions and preparation of Haematoxylin & Eosin (H&E)-stained slides used to develop the described tool were derived from previously completed experiments from parallel studies associated with the development of new snakebite therapeutics. Only the blinded images, representing diverse lesion sizes, colours and intensities, were selected as training set data for the VIDAL algorithm. All images used in its creation have been made available in https://github.com/laprade117/VIDAL-Experiments. Methodological details of the type of *in vivo* experiments performed and H&E-slide preparation can be found in Hall, *et al*^19^.

### Scoring of H&E-stained sections of venom-induced lesions

Brightfield images of the H&E-stained skin cross-section slides were taken with an Echo Revolve microscope (Settings: 10x magnification; LED: 100%; Brightness: 30; Contrast: 50; Colour balance: 50), with at least three images taken per skin cross-section. Dermonecrosis within each skin layer of each of the nine tissue samples was scored using methods outlined by Hall *et al*.^19^, from which an overall dermonecrosis score between 0 and 4 (0 signifying no dermonecrosis, and 4 signifying 100% dermonecrotic tissue) was determined for each sample. A more severe pattern of tissue damage was observed histologically in the dark-lesions as compared to the light lesions. Therefore, to take this difference into account, the mean dermonecrosis score of the dark-lesions (severe necrosis), was divided by that of the light lesions (less severe tissue damage) to calculate a ‘dark lesion severity adjustment score’, which was determined to be 2.019 (see details in the Results section).

### Printout sheet

To allow for standardised analysis of the dermonecrotic lesions, as well as to support the image analysis algorithms, we used the same A4 printout sheet as in our prior publication^32^ (c.f. Supplementary Figure 1 | A4 printout template to be used for dermonecrosis assays), which the tissue samples were placed on (Fig. S2,S3).

### Description of machine learning guided approach of quantifying dermonecrotic activity

We next trained a machine learning algorithm to automatically identify dermonecrotic lesions, adjust for lighting biases, scale the image, extract the dermonecrotic lesion area and discolouration, and calculate the DnUs. This was then implemented in a tool coined VIDAL (Fig. 1), for which we also prepared standard operating procedures (Fig. S4).

### White balancing, scaling, and segmentation

First, the input images were white balanced and scaled, as described in our prior publication^41^. Thereafter, to identify and segment the dermonecrotic lesions, we applied a deep learning method based on the U-Net architecture^33^, which we have previously also applied to snake venom induced haemorrhage identification ^32^. Changes from the original U-Net architecture include replacing the deconvolution layers in the expanding path with bilinear upsampling, followed by a 2×2 convolution, adding batch normalisation layers and using padding in each convolutional layer to preserve image dimensions ^43,44^.

Our dataset consisted of 193 training images taken (each with up to three lesions) with a Sony DSC W-800 camera. Each image contains 1-3 lesions displaying varying amounts of necrotic damage with both the light and dark necrotic regions in each image annotated. To limit annotator bias, each image was annotated by 4 different annotators resulting in 4 masks per image, each mask containing three classes (background/no lesion, light lesion, and dark lesion). For evaluating performance, we set aside 20% of the images at random as our held-out test set and performed 5-fold cross-validation on the remaining 80% of the images, evaluating model performance on the held-out test set. This process was repeated 5 times to avoid test-set bias.

At the time of training, the images were split into samples of size 256 × 256 pixels and fed into the model in batches of 32 samples. Batches were created such that each sample has a 50% chance of having a masked section of dermonecrotic tissue according to at least one annotator. The mask used for training was sampled from the set of annotators at random. Data augmentations include flips, rotations, noise, blurring, sharpening, distortions, brightness, contrast, hue, and saturation adjustments. They were selected to simulate the possible variation in both the lighting environment as well as account for different built-in post-processing implementations in different types of cameras.

The models were trained using the Adam optimizer and with a learning rate of 0.0001 for 180 epochs. We used a loss function based upon a combination of the Mathews correlation coefficient (MCC) and cross-entropy^45^.

### Calculation of dermonecrotic units and minimum dermonecrotic dose

Snake venom induced dermonecrosis primarily manifests itself in two distinct macroscopic appearances: 1. a severe dark dermonecrotic lesion, and 2. a less severe light region of local tissue damage. To compute the dermonecrotic severity of a given lesion, we quantify the area of both pathologies (1 and 2) and then combined them via a weighted sum using our *in vitro* determined (c.f. 2.4) weighting of 2.019 to 1 into DnUs.

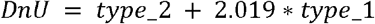

### Implementation in GUI

Using Streamlit (https://github.com/streamlit/streamlit) and localtunnel (https://github.com/localtunnel/localtunnel) with Google Colab, a simple web-based application to automatically analyse images was developed **(https://github.com/laprade117/VIDAL)**. A web-based application seems to be the most efficient way to quickly analyse data while working in the lab. Users can take a photo with a smartphone and upload it to the web-based tool (accessible via a smartphone browser) for an immediate result (Fig.S4).

## Acknowledgements

We would like to give our thanks to (i) Paul Rowley for maintaining the snakes at the LSTM herpetarium and for routine venom extractions, (ii) Dr. Laura-Oana Albulescu, Dr. Cassandra Modahl and Dr. Amy Marriott from LSTM for their help in planning and performing *in vivo* experiments, and (iii) Valerie Tilston and her team at the University of Liverpool for preparing the histopathology slides. The Authors acknowledge use of the Biomedical Services Unit provided by Liverpool Shared Research Facilities, Faculty of Health and Life Sciences, University of Liverpool. Funding was provided by a (i) Newton International Fellowship (NIF\R1\192161) from the Royal Society to SRH, (ii) a Wellcome Trust funded project grant (221712/Z/20/Z) to NRC, (iv) a UK Medical Research Council research grant (MR/S00016X/1) to NRC and (v) a UK Medical Research Council funded Confidence in Concept Award (MC_PC_15040) to NRC. This research was funded in part by the Wellcome Trust. For the purpose of open access, the authors have applied a CC BY public copyright licence to any Author Accepted Manuscript version arising from this submission.

## Author contributions

The project was conceived by TPJ, JMG, SRH, and WL. *In vivo* research was performed by KEB, EC, NRC, and SRH. The tool was designed by WL with input from TPJ, JMG, and SRH and architecture was checked by MM. Image annotations were performed by KEB, CRC, TDK, RNP, EC, CAD, and DSW and guided by JMG, SRH, WL and TPJ. JMG performed the histopathology scoring. The manuscript was drafted by TPJ and WL, with primary input from JMG, SRH, NRC, and AHL, as well as input from TJF, KEB, CRC, TDK, RNP, EC, CAD, DSW, MM.

## Data availability statement

The tool and sample images have been made available via GitHub (https://github.com/laprade117/VIDAL; https://github.com/laprade117/VIDAL-Experiments).

## Notes

https://github.com/laprade117/VIDAL

### Competing Interest Statement

The authors have declared no competing interest.

### Summary of Updates

Only the methods were rephrased to ensure clarity around the fact that no animal experiments were conducted for this study and rather data (i.e. images) from other studies involving such experiments were included.

