## Supplemental files for "Machine-learning guided Venom Induced Dermonecrosis Analysis tooL: VIDAL"

**Supplementary material**

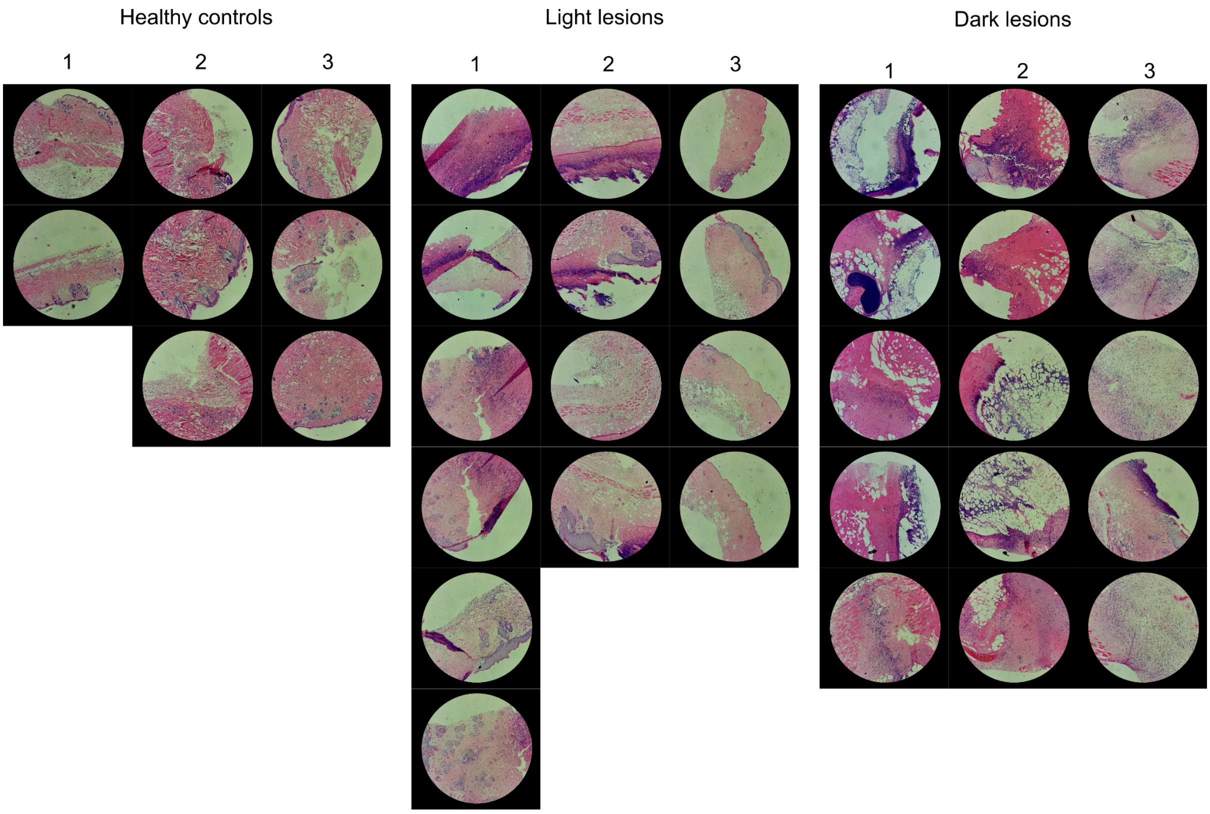


**Supplemental Fig. 1 All images of H&E-stained tissue samples used to quantify the %-dermonecrosis within each skin layer.** Images are 100x-magnified. Three tissue samples were chosen that exhibited either no necrosis (Healthy controls, left), large but light skin lesions (Light lesions, middle), and large and dark lesions (Dark lesions, right). Each number represents a different experimental animal.

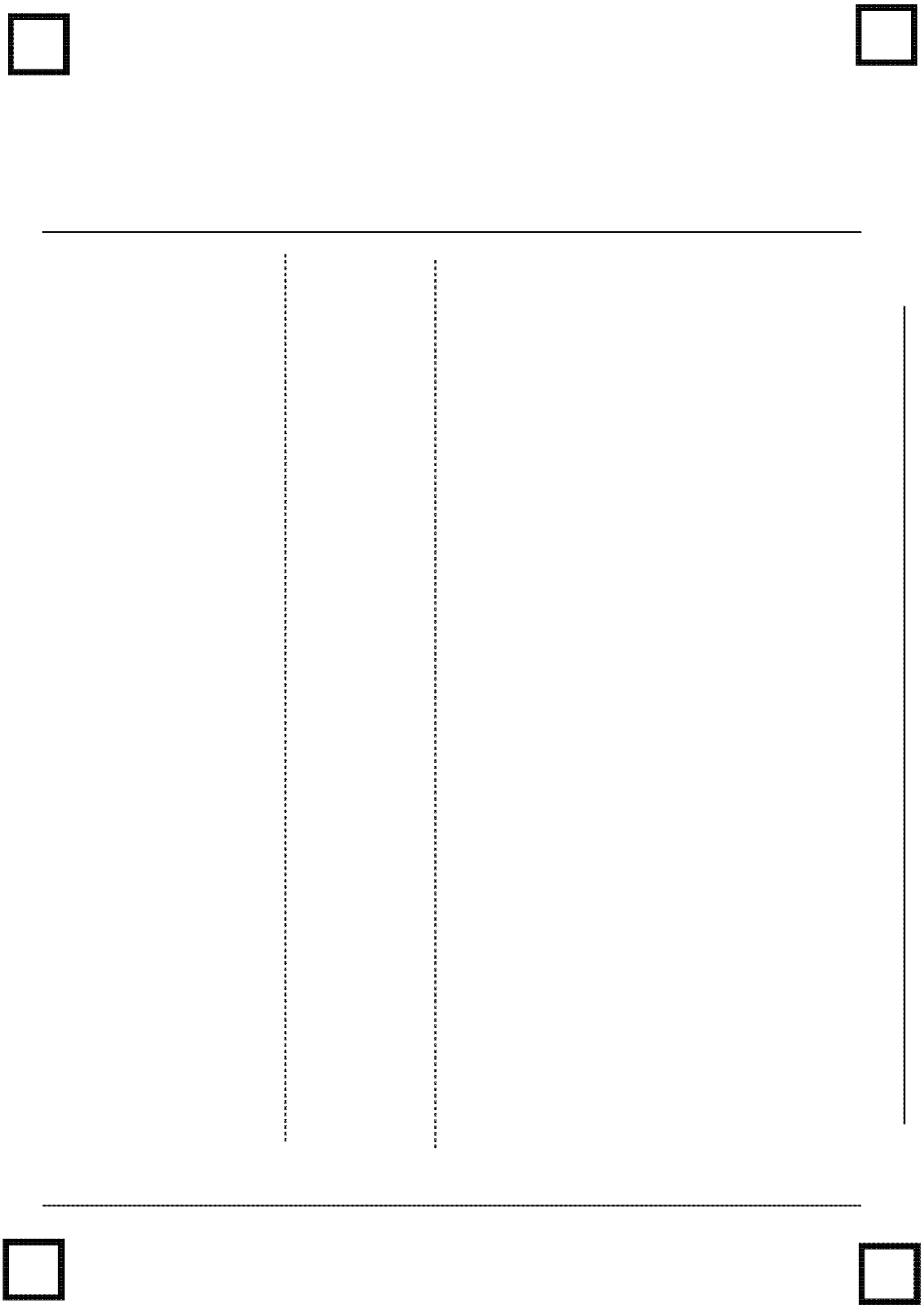

**Supplemental Fig. 2. A4 printout template to be used for dermonecrosis assays. This template is key, since it allows the tool to optimise scaling and better segment the lesions.**

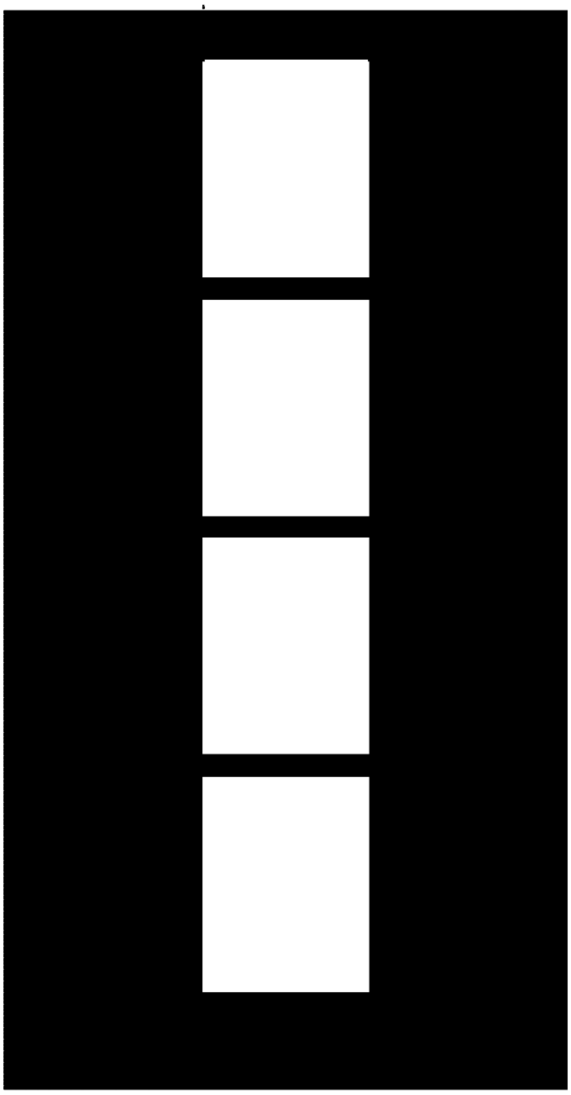

**Supplemental Fig. 3. A4 mask sheet to be used for dermonecrosis assays. This is to be placed on top of the mice, so each lesion is within one of the white cut-out boxes.**

**Supplemental Fig. 4. Standard operating procedure sheet for the VIDAL tool.**

### Machine-learning guided Venom Induced Dermonecrosis Analysis tooL: VIDAL

#### Standard Operating procedure

This document is meant to provide an easy-to-follow and step-by-step guide on how to perform dermonecrotic analyses using the VIDAL tool.

Materials

1. Print out template sheet (supplementary Figure 1)
2. Print out cover sheet (supplementary Figure 2)
3. Internet connection
4. Camera (e.g., phone)

Approach

1. To start with, place the euthanised mice or the excised lesion on the indicated lines of the template sheet (no more than four mice/lesions).
2. Place the black cover sheet on the mice/lesions, so that you have black boxes outlining the lesions. This helps the algorithm identify the lesions.
3. Ensure that all the black squares in each of the template sheet's corners are not covered and that you have as even lighting as possible. Aim to also minimise reflections on the lesions.
4. Take a photo of the sheet from above trying to avoid any tilt and so that each of the sheet's corners is equidistant to the camera.

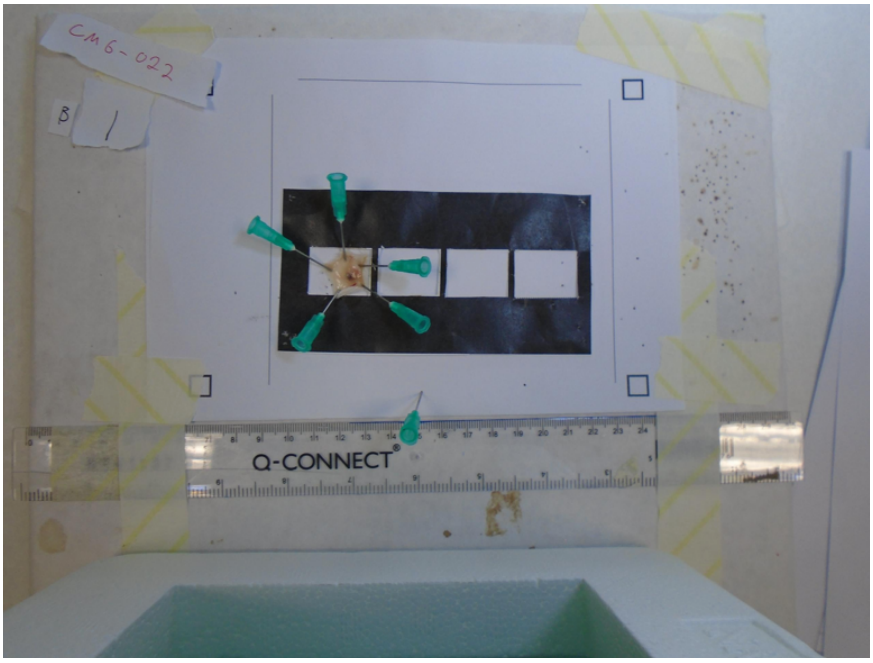

5. For ease of use it is recommended to transfer the image(s) to your computer, but you can also continue the analysis on your phone.
6. Open a browser window and follow the link to the tool: https://github.com/DTU-Bioengineering/VIDAL
7. Click the "Open in Colab" tab.

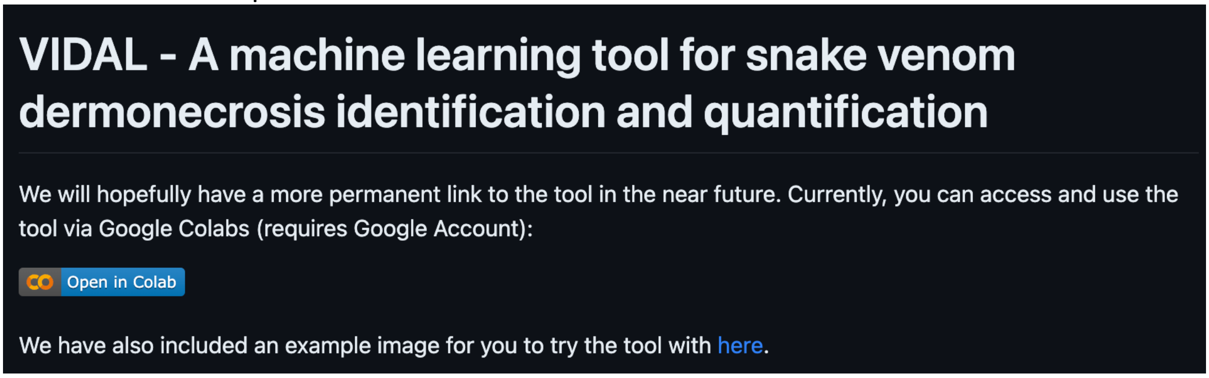


8. Now select "Runtime->Run all" and wait for each command cell (1-3) to finish running.

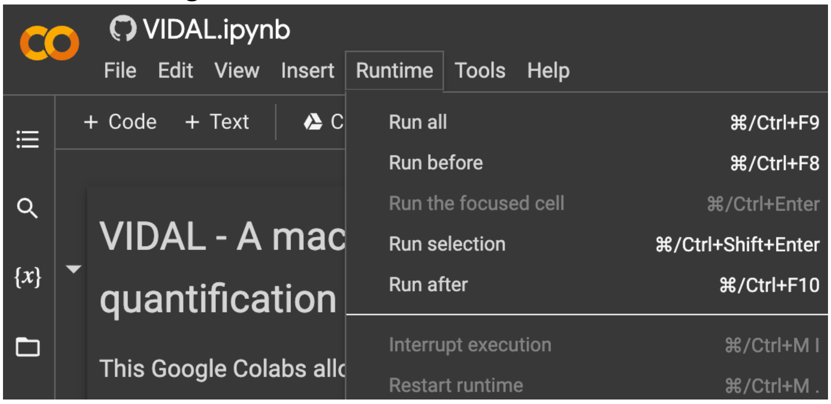


9. Once completed, click the url link the final line (e.g., https://aha-4030.loca.lt).

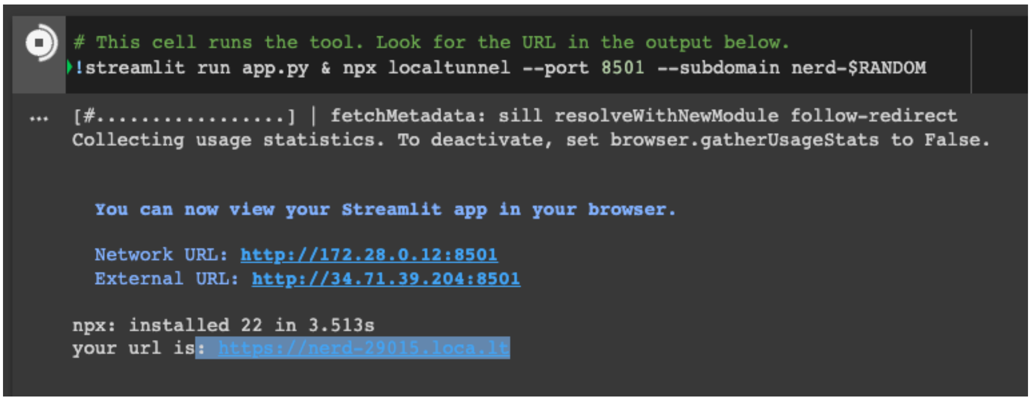


10. On the next page select "Click to continue"

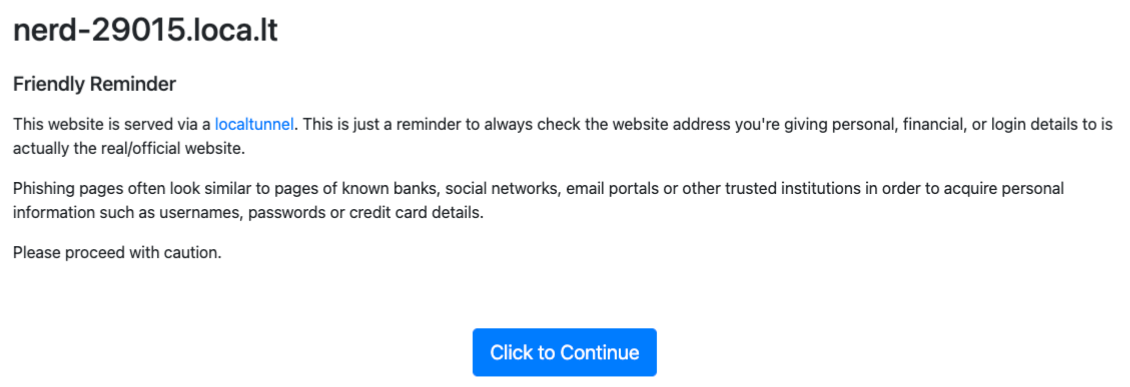


11. On the next page, upload your image as jpg, png, or tif (note: if your image is a jpeg instead of a jpg the tool might not recognise it. In that case just convert your image via one of the many free online image converters). You can also use the example image provided in the github folder, if you just want to try the tool.

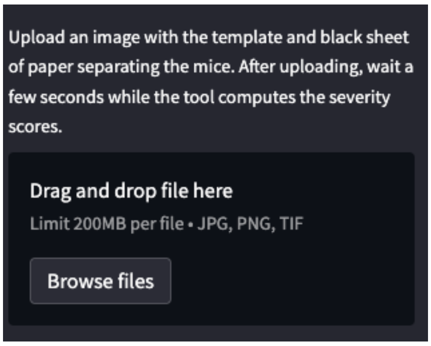


12. After uploading your image, wait a few sections you will be prompted to click on each of the lesions you want analysed (white dot). Afterwards just click compute.

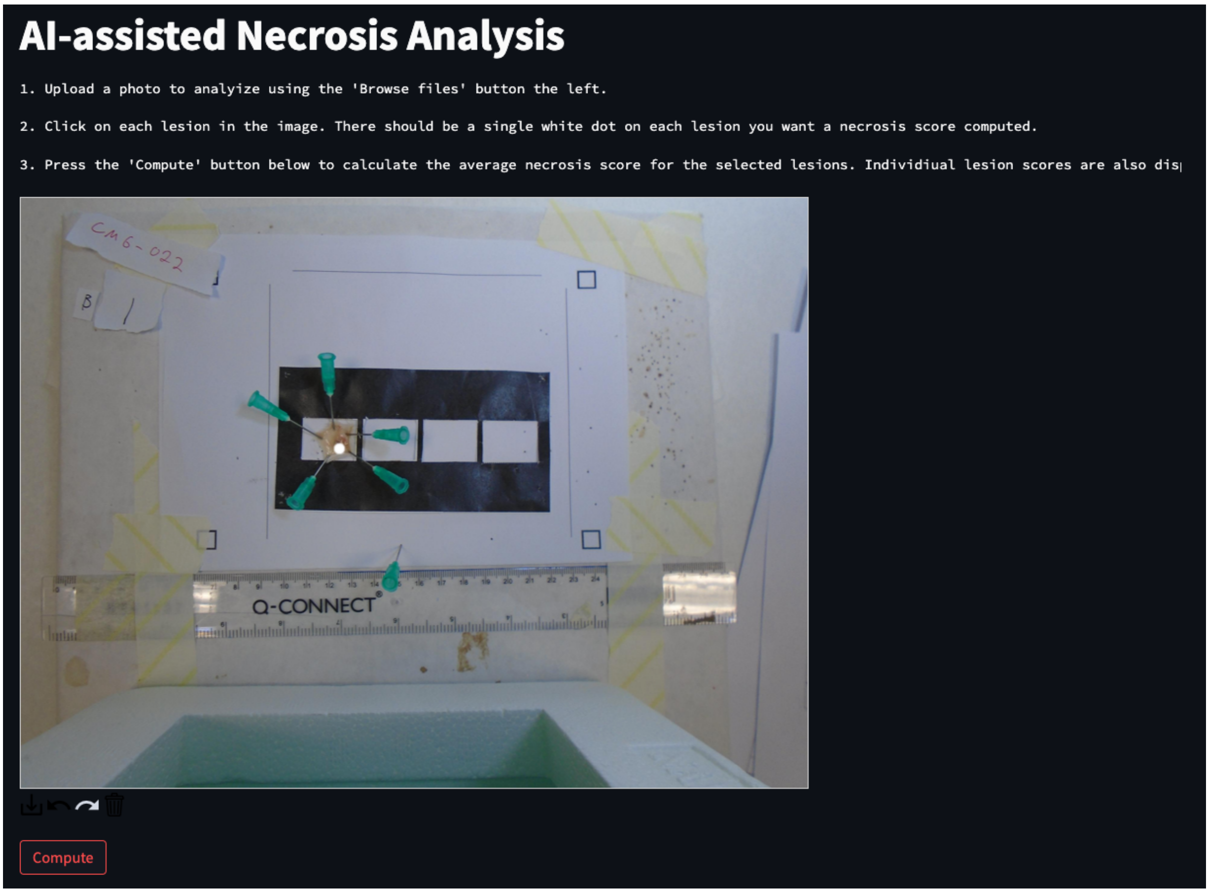


13. After a short time you should be presented with the following output, as well as a white balanced version of your image.

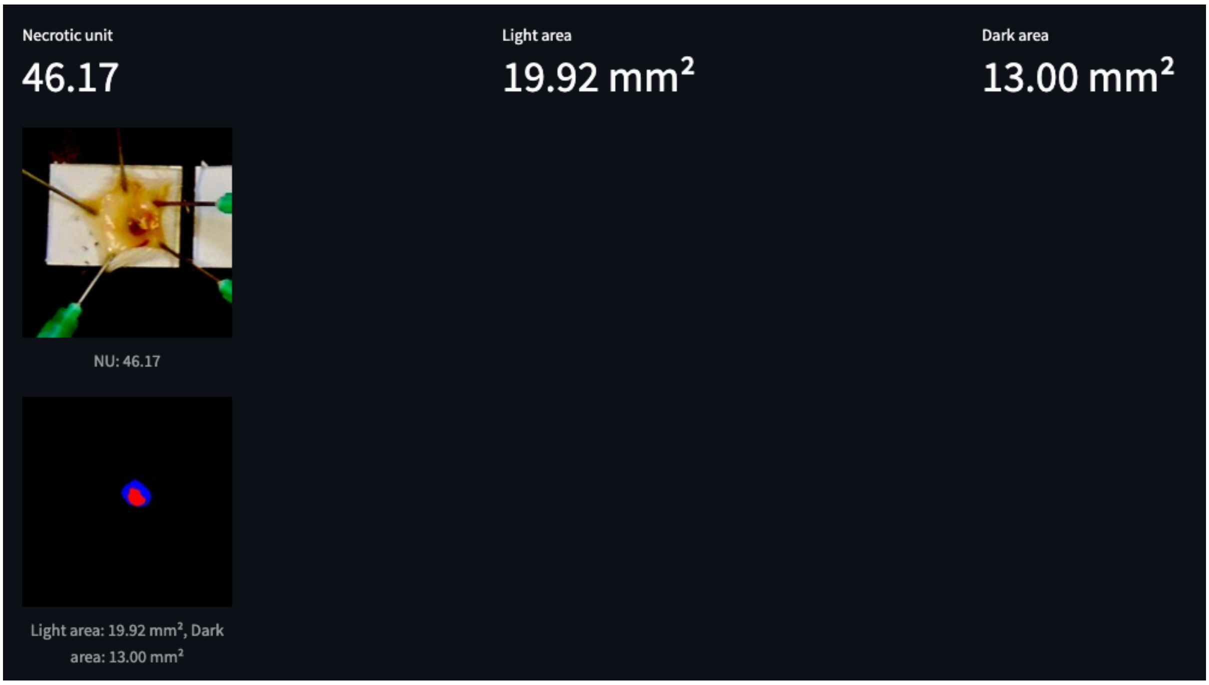
